# DigiLoCS: A Leap Forward in Predictive Organ-on-Chip Simulations

**DOI:** 10.1101/2024.03.28.587123

**Authors:** Manoja Rajalakshmi Aravindakshan, Chittaranjan Mandal, Alex Pothen, Christian Maass

## Abstract

Digital twins, driven by data and mathematical modelling, have emerged as powerful tools for simulating complex biological systems. In this work, we focus on modelling the clearance on a liver-on-chip as a digital twin that closely mimics the clearance functionality of the human liver. Our approach involves the creation of a compartmental physiological model of the liver using ordinary differential equations (ODEs) to estimate pharmacokinetic (PK) parameters related to on-chip liver clearance.

The objectives of this study were twofold: first, to predict human clearance values, and second, to propose a framework for bridging the gap between *in vitro* findings and their clinical relevance. The methodology integrated quantitative Organ-on-Chip (OoC) and cell-based assay analyses of drug depletion kinetics and is further enhanced by incorporating an OoC-digital twin model to simulate drug depletion kinetics in humans.

The *in vitro* liver clearance for 32 drugs was predicted using a digital-twin model of the liver-on-chip and *in vitro* to *in vivo* extrapolation (IVIVE) was assessed using time series PK data. Three ODEs in the model define the drug concentrations in media, interstitium and intracellular compartments based on biological, hardware, and physicochemical information. A key issue in determining liver clearance appears to be the insufficient drug concentration within the intracellular compartment. The digital twin establishes a connection between the hardware chip structure and an advanced mapping of the underlying biology, specifically focusing on the intracellular compartment.

Our modelling offers the following benefits: *i*) better prediction of intrinsic liver clearance of drugs compared to the state-of-the-art model and *ii*) explainability of behaviour based on physiological parameters. Finally, we illustrate the clinical significance of this approach by applying the findings to humans, utilising propranolol as a proof-of-concept example. This study stands out as the biggest cross-organ-on-chip platform investigation to date, systematically analysing and predicting human clearance values using data obtained from various *in vitro* liver-on-chip systems.

**Author summary:** Accurate prediction of *in vivo* clearance from *in vitro* data is important as inadequate understanding of the clearance of a compound can lead to unexpected and undesirable outcomes in clinical trials, ranging from underdosing to toxicity. Physiologically based pharmacokinetic (PBPK) model estimation of liver clearance is explored. The aim is to develop digital twins capable of determining better predictions of clinical outcomes, ultimately reducing the time, cost, and patient burden associated with drug development. Various hepatic *in vitro* systems are compared and their effectiveness for predicting human clearance is investigated. The developed tool, DigiLoCs, focuses explicitly on accurately describing complex biological processes within liver-chip systems. ODE-constrained optimisation is applied to estimate the clearance of compounds. DigiLoCs enable differentiation between active biological processes (metabolism) and passive processes (permeability and partitioning) by incorporating detailed information on compound-specific characteristics and hardware-specific data. These findings signify a significant stride towards more accurate and efficient drug development methodologies.

## 1. Introduction

The drug testing dilemma presents a significant challenge in pharmaceutical development, marked by high costs and a distressing attrition rate in accurately predicting human responses [1, 2]. A pivotal element in preclinical drug development is the accurate estimation of the first-in-human dose and different dosing regimens to keep drug levels within a therapeutic range. This demands precise assessments of hepatic clearance and human pharmacokinetics [3, 4]. Typically, the gold standard in drug development is the use of simpler *in vitro* systems to study drug metabolism, including liver microsomes [5] and suspension or plated hepatocytes [6]. The drug depletion data (time-concentration profile) are then analysed to determine the *in vitro* clearance rate. A very simple mathematical model is employed that considers the *in vitro* system as a single compartment, the one-compartment PK model [7]. Well-mixing and instantaneous drug distribution is assumed, all biological processes, e.g., permeability and partitioning from cell culture media into intracellular milieu are lumped into drug clearance. This approach also cannot differentiate between compounds actually being metabolised and compounds bound to media proteins or hardware. The so determined *in vitro* clearance value is then extrapolated to humans (*in vitro* – *in vivo* extrapolation) and integrated into human physiologically-based pharmacokinetic (PBPK) models [4, 8] to predict human pharmacokinetics (absorption, distribution, metabolism and excretion (ADME)), before actually testing a new compound in humans. Although this approach is well-established in drug development and easy to use, it also systematically underpredicts human PK [9] by 5-10 fold across studies and compounds.

Microphysiological systems (MPS) and organ-on-chips as well as 3D organoids hold great promise to address more complex *in vitro* ADME, toxicology and pharmacology questions offering miniature, biomimetic systems that replicate key aspects of human organ physiology [10, 11, 12]. These technologies create an environment where human cells can grow and interact in an organ-specific context, providing insights into human biology and disease that were previously unattainable in conventional *in vitro* models or animal studies. OoC and MPS-based systems are already used in today’s drug development for PK, but also for assessing drug efficacy and toxicity [13, 14, 10, 15]. While the emulated *in vitro* biology of MPS and OoCs is getting ever more complex and produces more human-relevant data, these systems still fall short in considerably improving the prediction power of in-human situations, like PK [16, 13]. However, MPS and OoC data are also still analysed using the state-of-the-art mathematical model (one-compartment), that is not accounting for the advanced biology. It remains unclear whether the OoC and MPS biology is still not human-relevant enough (and thus producing human-relevant data) or the state-of-the-art mathematical analysis is the cause for the underprediction. Potentially, a digital twin framework that enables the mapping of complex on-chip biology to advanced mathematical models could provide a useful approach to enable OoC and MPS translation to humans and increase the prediction power but is currently lacking.

The current study aimed at developing a digital twin approach integrating MPS and OoC data within advanced computational models of biology to improve the prediction of clinical clearances. DigiLoCs, our developed digital liver-chip simulator, facilitates the accurate description of on-chip complex biology. The tool comprises and utilises information on complex biological processes (clearance, permeability, partitioning), hardware-specific information from the studied *in vitro* system, and compound-specific information. By accounting for more multi-dimensional information, the tool enables differentiation between active biological processes, such as metabolism, and passive ones, such as permeability and partitioning of a compound from cell culture media into the cellular milieu. This contrasts with state-of-the-art approaches, where passive biological processes are not considered specifically and lumped together into a single process, i.e., clearance. Drug depletion kinetics of 32 compounds were taken from literature covering commercially available liver-on-chips (CnBio [13, 17], Javelin), and 3D spheroids [18, 19], including fast and slow-cleared compounds. According to these studies, DigiLoCs outperform the state-of-the-art prediction approach considerably. The impact of a more accurate description of clinical clearance values on predicting human PK was investigated in a proof-of-concept study using propranolol. The kinetics of propranolol was predicted in humans using the state-of-the-art, DigiLoCs, and literature approach. The results obtained from DigiLoCs for propranolol in the proof-of-concept study were much closer to the actual observed human values as compared to other approaches.

To the best knowledge of the authors, this is the first and biggest study so far, comparing head-to-head the performance of different hepatic *in vitro* systems to predict human clearance and demonstrating the impact OoC and MPS systems can have, in the drug development process enhanced through the modelling and prediction features of DigiLoCs.

## 2. Methods

In this section, we describe the following: *i*) data used in the study for predicting human clearance, *ii*) DigiLocs, digital twin for liver-on-chip, *iii*) mathematical model, parameter estimation and sensitivity analysis for DigiLocs, and *iv*) translation to humans and prediction of human pharmacokinetics.

### 2.1 Data

In this work, published data on pharmacokinetics (metabolism) or toxicology studies of 32 drugs are used (See Table 1) to predict human pharmacokinetics.

**Table 1.** Overview of literature reports providing on-chip pharmacokinetic information on compound clearance.

| Study | Drugs | Used in<br>this<br>study | *In vitro* system | cell<br>number<br>[a.u.] | Media<br>volume<br>[ml] | Flow | Compartments |
| --- | --- | --- | --- | --- | --- | --- | --- |
| Docci et al. 2022 [13] | 9 | 9 | CnBio Liver-Chip | 3E5 | 1.60 | Recirculation | 1 |
| Tsamandouras et al. 2017 [17] | 6 | 3 | CnBio Liver-Chip | 3E5 | 1.60 | Recirculation | 1 |
| Rajan et al. 2023 [20] | 12 | 8 | Javelin Liver-Chip | 2.15E5 | 1.30 | Recirculation | 2 |
| Kanebratt et al. 2021 [18] | 4 | 4 | 3D Spheroid (Hurel) | 6E3 | 0.05 | No | 1 |
| Bonn et al. 2016 [19] | 8 | 8 | 3D Spheroid (Hurel) | 3E4 | 0.10 | No | 1 |
| Total | 38 | 32 |  |  |  |  |  |

### 2.2 DigiLoCs: Digital Twins for cell-based liver assays

DigiLoCs is a software tool that describes the on-chip complex biology more accurately in the context of use to predict clinical clearance values. The software comprises (Fig 1):

**Fig 1.**
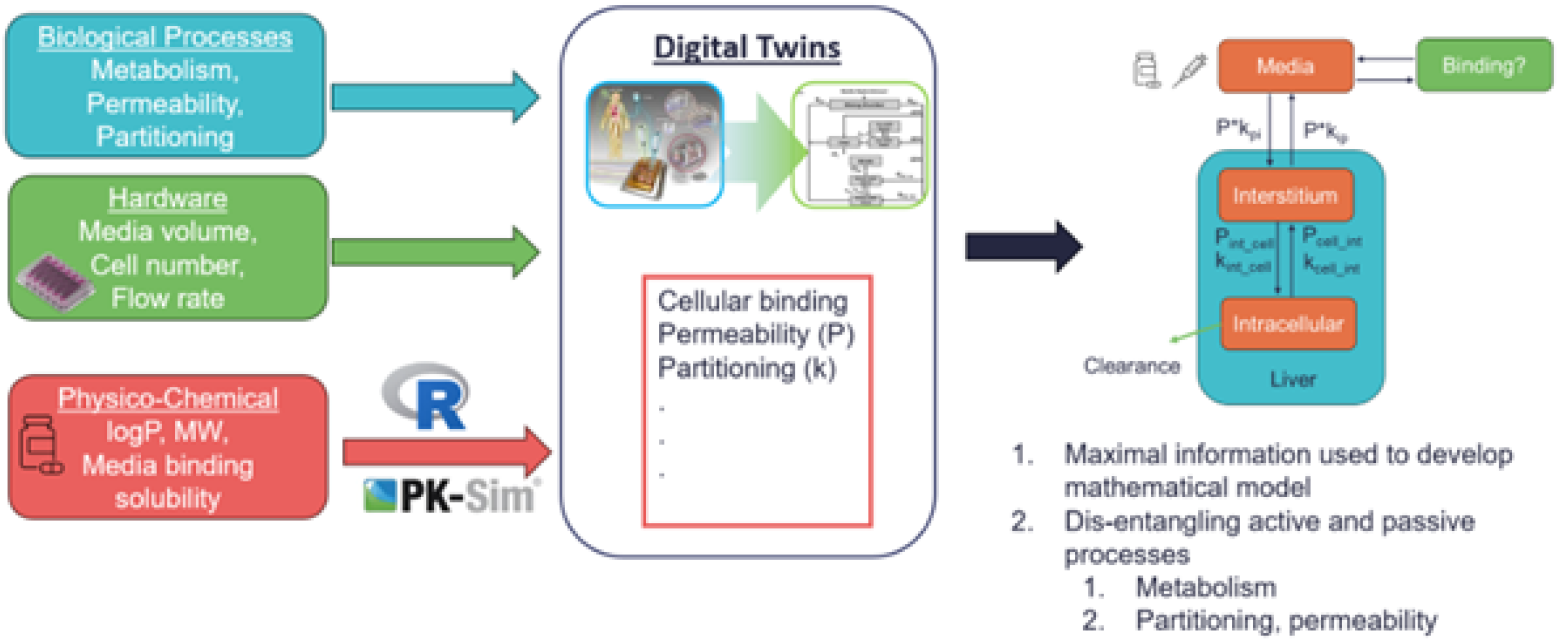
Digital Twin (DT) Approach. Contrasting state-of-the-art, the DT approach uses biological, hardware, and physicochemical information to map the biological processes on-chip more accurately to *in silico*, thereby maximising the information leveraged. This results in the disentanglement of active (metabolism) and passive (permeability, partitioning) processes.

- modelling of complex biological processes (clearance, permeability, partitioning),
- hardware-specific information from the studied *in vitro* system and
- compound-specific information.

The tool differentiates between active biological processes, such as metabolism, and passive ones, like permeability and partitioning of a compound from cell culture media into the cellular milieu. This contrasts with state-of-the-art approaches, where passive biological processes are not considered especially and lumped together into a single process, i.e., clearance.

Liver-on-chip technology provides a more physiologically relevant environment compared to traditional cell cultures or animal models, enhancing the simulation of drug responses using mathematical models. Hence, a more accurate mathematical description is needed. The three primary compartments of the liver chip considered in the model are media, interstitium, and intracellular space, which serve as dynamic environments where drugs are distributed, metabolised, and interact with hepatic cells. This compartmentalisation is based on concepts applied in human whole-body PBPK modelling.

The software tool is developed in the open-source programming language R and seamlessly communicates with PK-Sim (https://www.open-systems-pharmacology.org/) via in-house developed functions. For more information, see esqlabsR package (https://github.com/esqLABS/esqlabsR). All analysis and plotting were done in the open-source programming language R. Moreover, the proposed workflow does not interfere with existing wet lab Standard Operating Procedures (SOPs) for performing biological experiments and does not add an extra considerable burden to the user. DigiLoCs uses existing biological data, and its performance may be improved by measuring cell-associated compound concentrations in addition to the compound media depletion time course, which would add a minor extra step in the lab SOP. This, however, is negligible given the improvement in performance power and the confidence in the prediction.

#### 2.2.1 Implementation of hardware specifications

DigiLoCs map the chip architecture to a compartmental model to describe the time-dependent distribution of a compound on-chip. The compartment models use time-dependent ordinary differential equations (ODEs) and assume well-mixing within compartments. These are generally accepted to describe the distribution of exogenous and endogenous compounds and molecules.

A physical chamber separated by a membrane or connected by flow to another chamber is represented by a compartment in the software. Serial compartments are connected via concentration-dependent flow rates (typically in *ml/min*) between the compartments and normalised by the volume of the originating compartment.

#### 2.2.2 Implementation of biological specifications

The biology (more precisely, the cell type exerting the biological function under investigation; here: metabolism) is mapped by two additional compartments representing the interstitial and intracellular space of the investigated biology (Fig 1). Transition rates from the cell culture media into the interstitial and intracellular milieu are described by two core processes:

- permeability (*cm/min*, how fast is a compound taken up?)
- partitioning (how much of the compound is taken up by cells?)

These processes are proportional to the time-dependent concentration of the compound and the surface area shared between the channels and the cell layer. Lastly, the metabolism rate is allocated at the intracellular compartment and corrected by the unbound fraction of the compound in the intracellular compartment. Similar to the compartmental structure are these core processes accepted in describing the distribution of compounds in a pharmacokinetic framework.

#### 2.2.3 Implementation of compound-specific information

The following physicochemical properties of the investigated compounds are used in the software: *i*) lipophilicity (logP), *ii*) molecular weight (MW) and *iii*) fraction unbound (fu); to calculate up to six dependent downstream parameters (listed below). These parameters describe the partitioning from the main media compartment into the interstitial space and between the water fraction and both interstitial and intracellular space. Additionally, permeability across the endothelial barrier and between interstitial and intracellular spaces is calculated.

Partitioning: *i*) K_int___pls_ *ii*) K_water _ cell_ *iii*) K_water___int_

Permeability: *i*) P_endothelial_ *ii*) PA_cell___int_ *iii*) PA_int___cell_

Here *int* refers to interstitial space, *pls* refers to media, *water* refers to water exchange fraction, and *cell* refers to intracellular space. These parameters are calculated based on well-established and documented equations implemented in PK-Sim [3]. The same partition coefficient calculation methods as implemented in PK-Sim are also readily available and can be investigated:

- PK-Sim standard
- Poulin and Theil
- Rodgers and Rowland
- Schmitt
- Berezhkovskiy

Further, only the unbound fraction of a compound can be taken up by cells and be metabolised by cells. The unbound fraction in the cell culture media is typically informed by biological experiments. However, the intracellular unbound fraction is not often available or measured. Thus, two established QSAR models (quantitative structure-activity relationship) are implemented in the software to predict the unbound intracellular fraction of the investigated compound as a function of its physicochemical properties [21, 22].

### 2.3 Mathematical model

A typical single-compartment model is described as: Let *C*(*t*) be the drug concentration in the chip at time *t, V* the volume of the chip, and CL_*c*_ the clearance parameter. Then we have

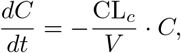

with initial value *C*(*t* = 0) = *C*_0_. The solution to this ODE is

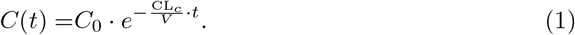

Taking the logarithm on both sides,

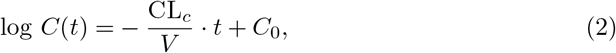

which is equivalent to regression on log-transformed kinetic data.

We developed a digital twin of liver-on-chip with three compartments that incorporates much more information on parameters related to both on-chip characteristics and drug-specific properties. The three-compartment model considering media, interstitium and intracellular compartment are described as follows: Let *C*_*p*_(*t*), *C*_*i*_(*t*), *C*_*c*_(*t*) and *V*_*p*_, *V*_*i*_, *V*_*c*_ be the concentration of the drug at time *t*, and volume of the plasma, interstitium, and intracellular compartment respectively. CL_*c*_ is the clearance parameter. Then we have

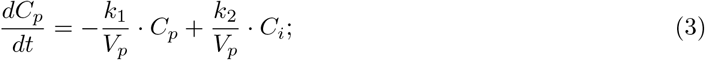

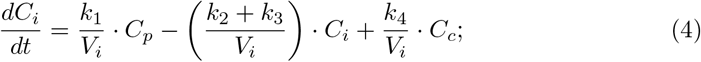

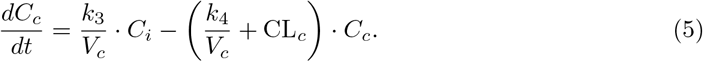

The parameters here are defined as follows,

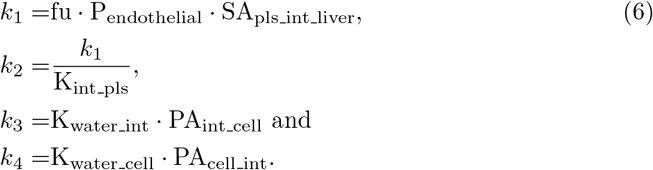

where K_int_pls_ is the rate constant for the transfer of drug between the plasma and interstitium, P_endothelial_ is the permeability coefficient of the drug between the endothelial layer, SA_pls_ int_ liver_ is the surface area of the interstitium, K_water int_ is the rate constant for water exchange or movement within the interstitium, K_water _cell_ is the rate constant for water exchange or movement within the intracellular compartment, PA_cell_ int_ is the permeability coefficient of the cellular membrane in the intracellular compartment, PA_int _cell_ is the permeability coefficient of the cellular membrane in the interstitium compartment and fu is the fraction unbound (plasma, reference value).

We can write this system of linear ODEs in matrix form,

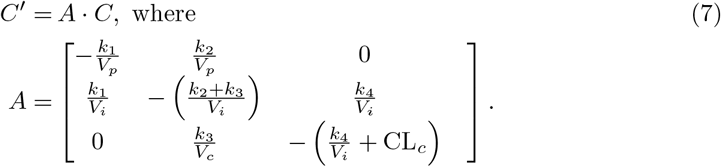

The general solution for the system of ODEs at time *t*:

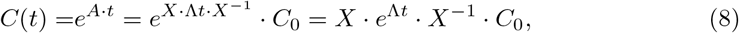

where *X* is the matrix of eigenvectors of *A*, Λ is the diagonal matrix with eigenvalues *λ*_1_, *λ*_2_, *λ*_3_ as diagonal entries, and *C*_0_ is the initial value of variables at time 0.

The objective function for optimisation is as follows:

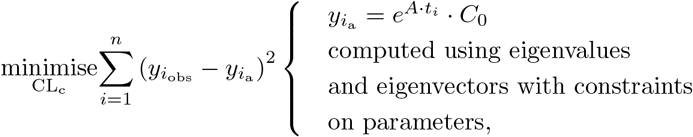

where 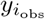 is the observed data point and 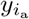 is the computed value using the eigenvalues and eigenvectors at the time *i* respectively, *n* is the number of observed data points.

### 2.4 Parameter Estimation

Parameter estimation aims to find unknown parameters in a computational model and is estimated using experimental data collected from well-defined and standard conditions. By minimising the distance of theoretical function values and experimentally known data, the set of parameters in the model can be estimated. The parameters which are not directly measurable can be estimated using least squares or any other fitting methods to analyse the model quantitatively. Nominal parameter values are obtained from PK-Sim, which is a comprehensive software tool for whole-body PBPK modelling. It enables rapid access to all relevant anatomical and physiological parameters for humans and common laboratory animals contained in the integrated database for model building and parameterisation.

Parameter estimation in DigiLoCs is a two-step process. Firstly, a customised cost function is implemented. This cost function calculates the weighted difference (ssq) between the model simulation (pred) from a specific compartment and the corresponding observed data (obs) for each time point according to:

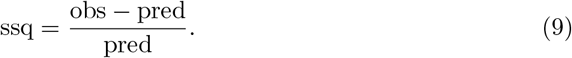

Common parameter estimation methods include maximum likelihood estimation and Nelder Mead optimisation. Nelder Mead, a non-linear optimisation method, is used to find the minima of the objective function in this work. Additionally, the partition coefficient between the intracellular (IC) and the main media compartment is estimated using the area under the simulated time-concentration profile of the IC and interstitial (IST) compartment and corrected for by the QSAR-predicted cellular unbound fraction (fu_cell_) and the unbound fraction in the media (measured, fu_media_):

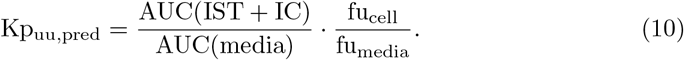

Kp_uu,obs_ is calculated from literature [23], where media and intracellular concentrations in hepatocytes were determined. Initially investigated for suspension hepatocytes in a 2D setting, the authors provide a scaling factor (*∼*4.9) to apply to human hepatocytes. Further, the ionisation state of the investigated compound (-1, 0, 1) results in a different partitioning. Otherwise, a range of possible partition coefficients are investigated. This is an additional anchor point for estimating the cost function value and links the simulated intracellular and main media compartment concentrations. Eventually, both differences are squared and summed up, resulting in the final sum of residuals. Based on this, a compound-specific scaling factor (SF) is calculated and used to scale the predicted human clearance:

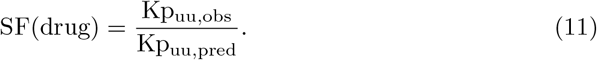

Specifically for the liver use cases, on-chip liver clearance and surface area between the main plasma compartment and the cell layer are estimated. It is possible to estimate other parameters, such as pre-calculated permeability or partition coefficient values.

#### 2.4.1 Implementation of software

Methodologically, DigiLoCs is implemented in the open-source programming environment R with its own package. A library of two common chip architectures and two cell types with six different chip-specific settings are already implemented:

1. One chamber, no media flow
2. Two chamber, recirculating flow
3. Organ-on-chip (hepatocytes)
  a. CnBio
  b. Hurel 1
  c. Hurel 2
  d. Dynamic42
  e. Javelin

These building blocks can be interchangeably used and connected, similarly to the building blocks in PK-Sim. While the R code and its package provide a step-by-step guide to generate and run a simulation, the code communicates seamlessly with a generic PK-Sim model to determine partitioning and permeability values as described above, which are used in the simulation.

### 2.5 Sensitivity Analysis

Both local and global sensitivity analyses are used to quantify the impact of input parameters on the output variables. This involves varying certain input parameters and observing changes in the output variable, intracellular concentration. The local sensitivity is estimated by changing one input parameter at a time while other parameters are held constant. The study provides insights into the sensitivity of various parameters and how they affect the output of the model.

The input parameters K_int _pls_, P_endothelial_, SA_pls_int_liver_, K_water_ int_, K_water_cell_, PA_cell _int_, PA_int _cell_, fu and CL_*c*_ are varied to evaluate the sensitivity of the output variable intracellular concentration (*C*_*c*_). First, the local sensitivity is estimated by changing one input parameter by 10% at a time while the other parameters are held constant, and the changes in output variable *C*_*c*_ are compared. Eq 12 shows the local sensitivity index for *C*_*c*_ with respect to the varying model parameter (P_*i*_), which is approximated by a small perturbation ΔP_*i*_,

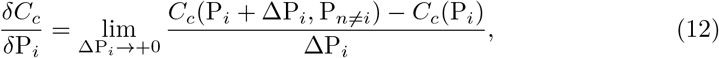

where *C*_*c*_(P) is the model prediction of the intracellular concentration for parameter set P. The local sensitivity index is normalised to eliminate the effect of units:

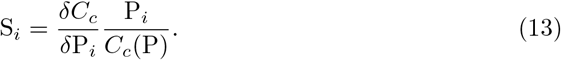

Global sensitivity analysis evaluates the effect of potential interactions of the input parameters in an output variable. The Sobol sensitivity analysis of the SALib package in Python is used to perform the global sensitivity analysis. Input parameters are sampled using the Saltelli sampler. The lower and upper bound of the parameters are set as 0.1-fold and 10-fold of the baseline parameter values, respectively. The first-order and total-order indices are estimated using the Sobol sensitivity analysis.

### 2.6 Translation to Humans

Drug-related parameters extracted from OoC or any other *in vitro* studies can be scaled to predict clinical parameters using *in vitro*-*in vivo* translation (IVIVT) [17, 13]. The typical value of unbound intrinsic clearance CL_int(u)_ determined for each drug from the pharmacokinetic analysis of the *in vitro* depletion data is scaled up to a human liver equivalent unbound intrinsic clearance CL_int(u),H_ using

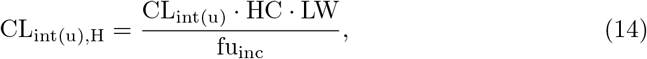

where HC is the human hepatocellularity of 120 million cells / g of liver, LW is the average human liver weight of 25.7g / kg of body weight [17] and fu_inc_ is the unbound fraction of drug in the incubation medium. The hepatic clearance (referring to whole blood concentrations) is then predicted (CL_H,pred_) using the Well-Stirred (WS) model:

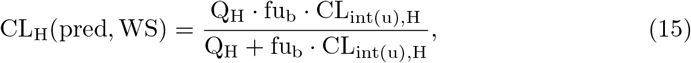

where Q_H_ the average hepatic blood flow of 20.7 mL/min/kg of body weight and fu_b_ is the fraction of the drug unbound in blood. The fraction unbound in the blood (fu_b_) was calculated for each compound from the known fraction unbound in the plasma (fu_p_) and blood-to-plasma ratio (Rbp) according to the equation fu_b_ = fu_p_*/*Rbp or directly used, if available from the literature. The predicted hepatic clearance, CL_H,pred_ values were then compared to observed hepatic clearance, CL_H,obs_ values (referring to whole blood concentrations).

### 2.7 Prediction of Human Pharmacokinetics

First, a physiologically-based kinetic model (PBK) is developed using qualified installations of the PBK software PK-Sim. A whole-body PBK model includes an explicit representation of the organs most relevant to the uptake, distribution, excretion, and metabolism of the drug. These typically include the heart, lungs, brain, stomach, spleen, pancreas, intestine, liver, kidney, gonads, thymus, adipose tissue, muscles, bones, and skin. More information can be found in S1 Text.

The tissues are interconnected by arterial and venous blood compartments, and each is characterised by an associated blood flow rate, volume, tissue partition coefficient, and permeability. If applicable, R (Distribution 4.0) and RStudio (Version 1.2.5) are used in the analysis for preprocessing and post-processing of data and model outputs [24]. The analytical approach is based on the principles set out in the guidelines of the EMA, FDA, and/or OECD for reporting on PBK M&S [11]. The developed PBK model is used to describe the human kinetics of propranolol. Key kinetic parameters are informed by either clinical data, literature values or on-chip predictions. The translational workflow that integrates organ-on-chip results to predict human pharmacokinetics is shown in Fig 2.

**Fig 2.**
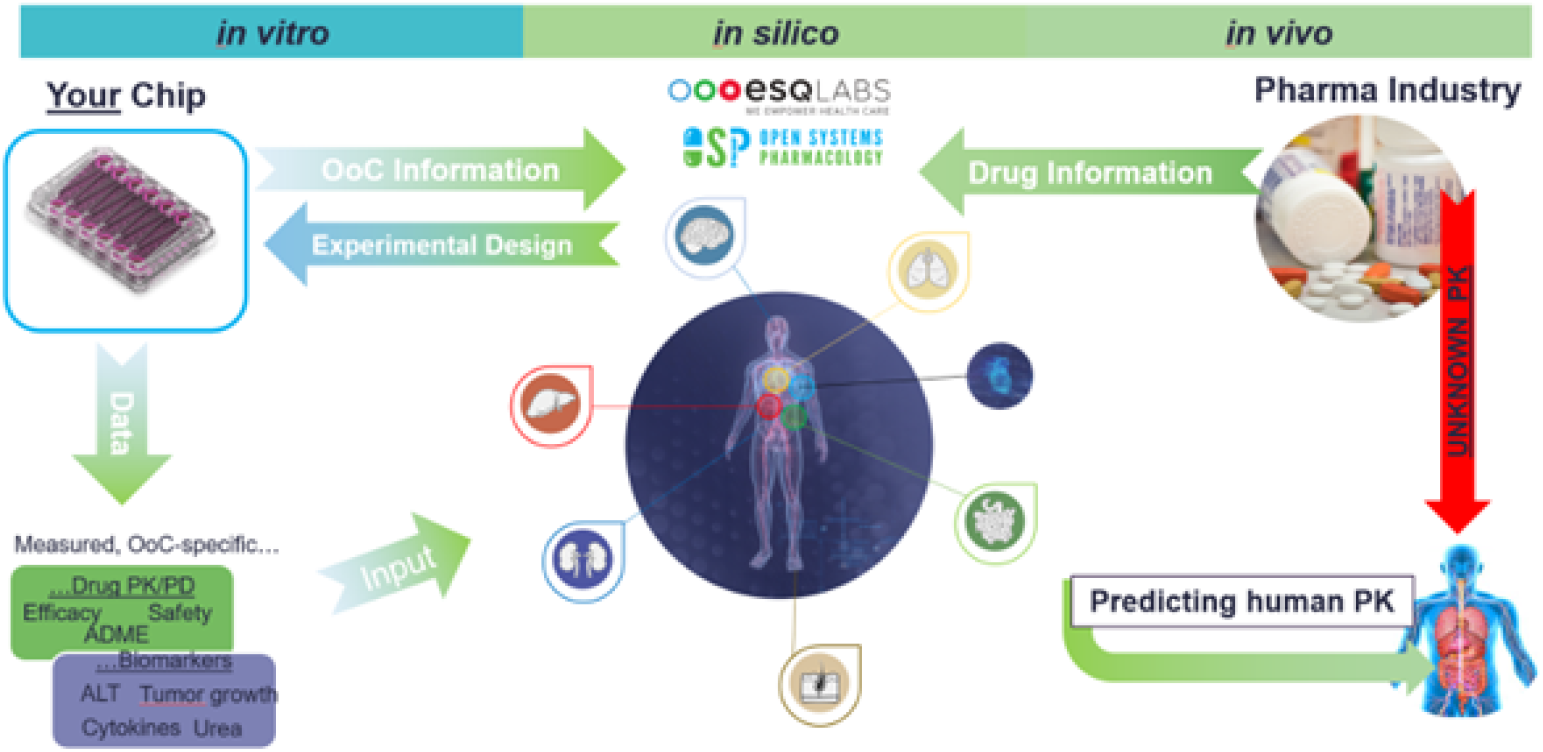
Translational workflow plan that integrates results from organ-on-chip with computer modelling to predict the kinetics of drugs in humans. The digital twins of the humanised organ-on-chip systems together with chip-specific information and physicochemical information are developed in R.

## 3. Results

The liver clearance and surface area of the chip are estimated after fitting the drug kinetic data. This section describes the following: digital twin-based model simulations for selected compounds, sensitivity analysis results, predicting human clearance, and translation to human PK using propranolol as a proof-of-concept study. The Poulin and Theil method of partition coefficient calculation was used due to its superior fit to observed drug kinetics, as evidenced by lower residual error (data not shown), outperforming alternative methods.

### 3.1 Simulating compound depletion on-chip

The digital twins for the investigated *in vitro* liver systems were successfully implemented in R and used to simulate the on-chip kinetics. After parameter estimation, the resulting model simulations describing the observed compound depletion data were visually inspected. The final parameter values can be found in Tables A and B in S1 Text. Additionally, the squared sum of residuals was evaluated and deemed acceptable *<* 0.01, which was the case for all simulations (data not shown). An example of on-chip kinetics is presented in Fig 3. As can be seen, the digital twin approach (violet line) captures the on-chip kinetics (blue dots) very well. Simultaneously, the intracellular (IC) kinetics are plotted (red lines), clearly highlighting the difference in compound uptake and, thus clearance rates. The remaining figures are presented in S1 Text (See Figs A-D).

**Fig 3.**
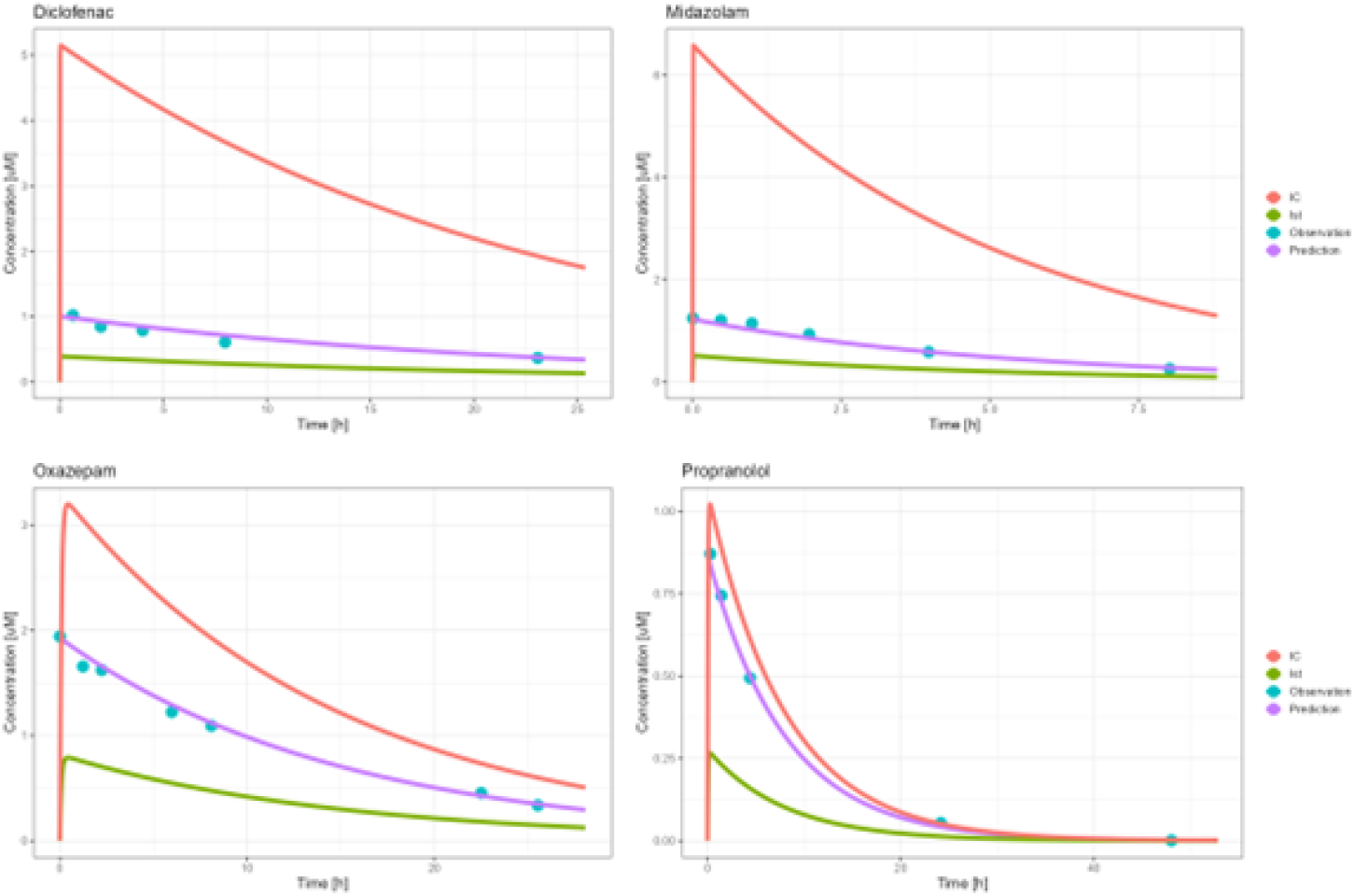
Digital twin-based model simulation of on-chip kinetics after fitting parameters for selected compounds; diclofenac, midazolam, and oxazepam are from Docci et al. [13], while propranolol is from Tsamandouras et al. [17]; IC = intracellular, Ist = interstitium.

### 3.2 Sensitivity analysis

The sensitivity analysis, both local and global, was conducted to quantify the sensitivity of model output intracellular concentration with input parameters. The analyses were performed for various parameters, and the results indicated that the output is more sensitive to parameters such as the permeability coefficient of the endothelial layer, surface area of the liver sinusoids, and clearance.

These parameters were estimated or calculated from experimental results. Clearance (CL_*c*_) is identified as the most sensitive parameter with respect to intracellular concentration. The results imply that accurate values of these sensitive parameters are crucial for the model’s accuracy.

The normalized local sensitivity indices (Fig 4a) and the first-order and total-order global sensitivity indices (Fig 4b) for intracellular concentration across the input the parameter set is shown. The results from both local and global sensitivity analyses shows that the output is more sensitive to parameters P_endothelial_, SA_pls int liver_, fu and CL_*c*_. These parameters were estimated or calculated from experimental results. Clearance (CL_*c*_) is the most sensitive parameter with intracellular concentration. SA_pls int liver_ and CL_*c*_ were estimated and nominal values were used for all other parameters. SA results imply that we need correct values of the constants P_endothelial_, fu as they are more sensitive.

**Fig 4.**
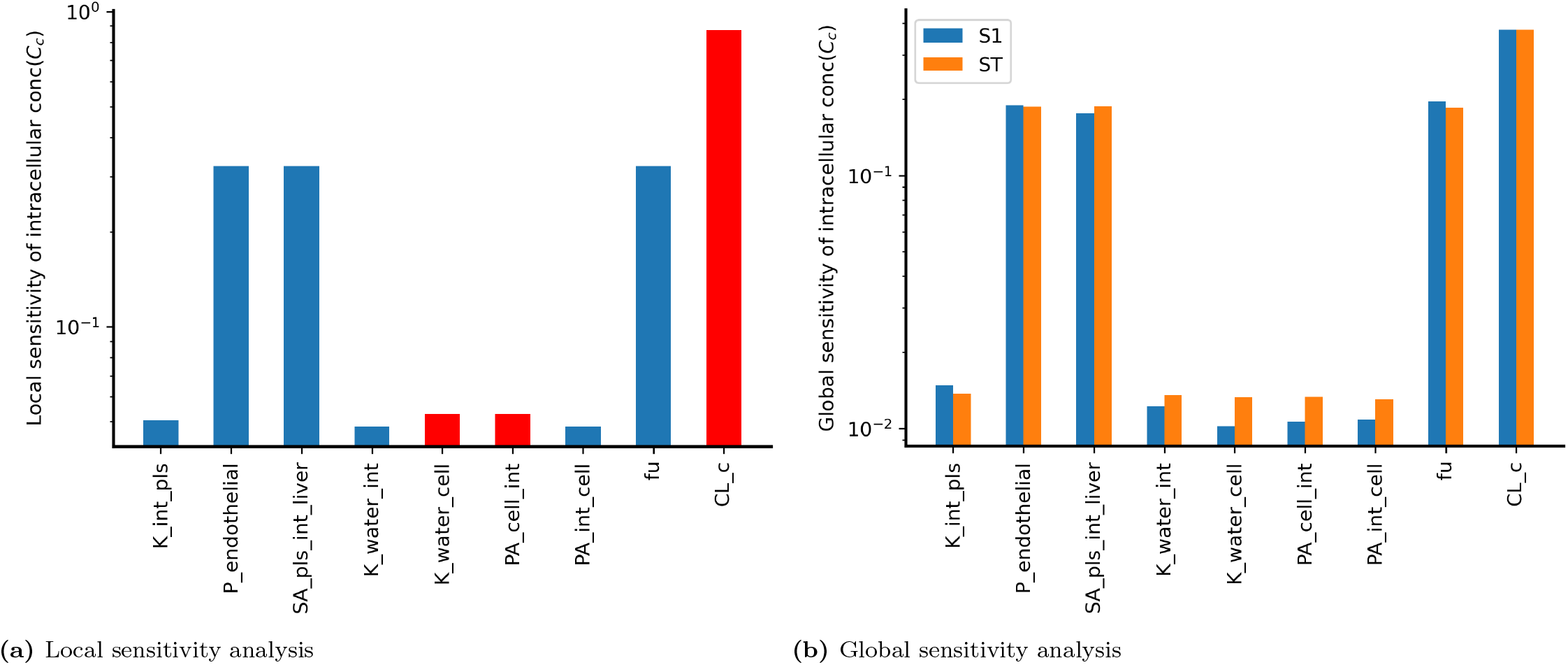
Local and global sensitivity of the parameters with respect to output intracellular concentration. (a) Blue bars indicate that the output and the input change in the same direction and the red bar indicates that the output decreases when the input increases. (b) The blue and orange bars represent first-order and total-order indices, respectively.

### 3.3 Predicting Human Clearance

Following, the on-chip estimated clearance values were translated to total human clearance according to Eq 15. More detailed information is available in S1 Text (see Table A and B). Likewise, from the investigated studies (Table 1), *in vitro* unbound clearance values were available and scaled to human equivalents.

Following, the ratio of clinical observed human clearance values and either predicted human clearances using state-of-the-art mathematical modelling or the digital twin approach were estimated and converted into a density function for easier graphical visualisation. As can be seen in Fig 5, the digital twin approach (DigiLoCs) outperforms the standard approach considerably. The center of the distribution is around 1 indicating a non-biased prediction of clinical clearance values, while the width of the distribution is very small. Quantitatively, the ratio for the digital twin approach over all compounds is 1.04 *±* 0.31, with a coefficient of variation of 30%. In contrast, the standard approach (blue curve) majorly under-predicts the clearance values while maintaining a broad distribution and thereby adding to uncertainty in the prediction (0.56 *±* 0.44, CV = 79.3%). The correlation plot between the observed and herein predicted clinical clearance values highlights on a drug-individual level the improved prediction performance of the DigiLoCs approach. As can be seen in Fig 6 (example graph for CnBio Liver-on-Chip data), most of the compounds fall within the 1.5-fold line (Average fold error, AFE = 0.965). Similar correlation plots are presented in the S1 Text (See Figs E-G) for the other *in vitro* systems.

**Fig 5.**
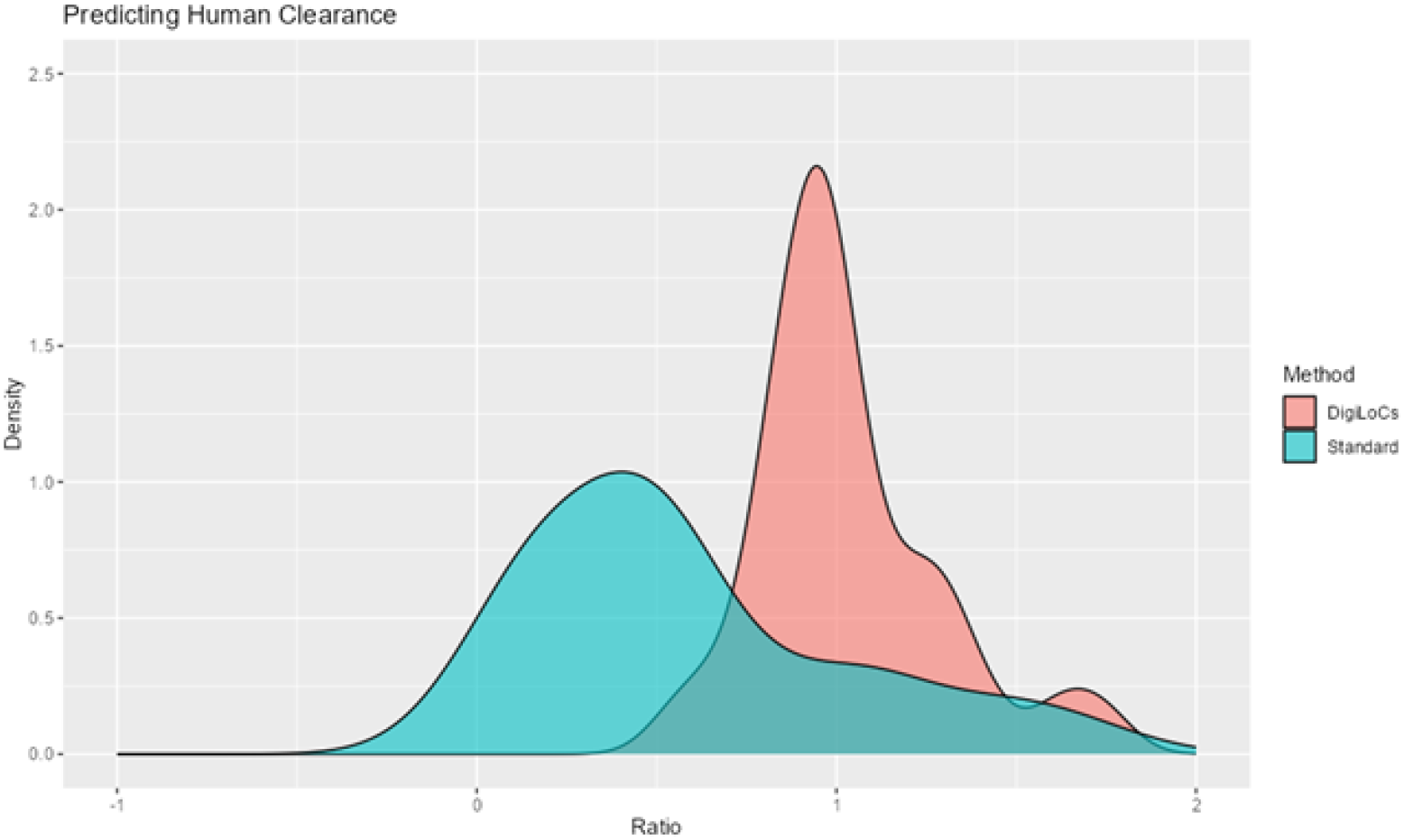
Impact of DigiLoCs on predicting clinical clearance values compared to the state-of-the-art approach. In total, a set of 32 compounds across three different *in vitro* liver-systems have been investigated. The x-axis presents the ratio of predicted/observed clinical clearance values using either the DigiLoCs or the state-of-the-art approach.

**Fig 6.**
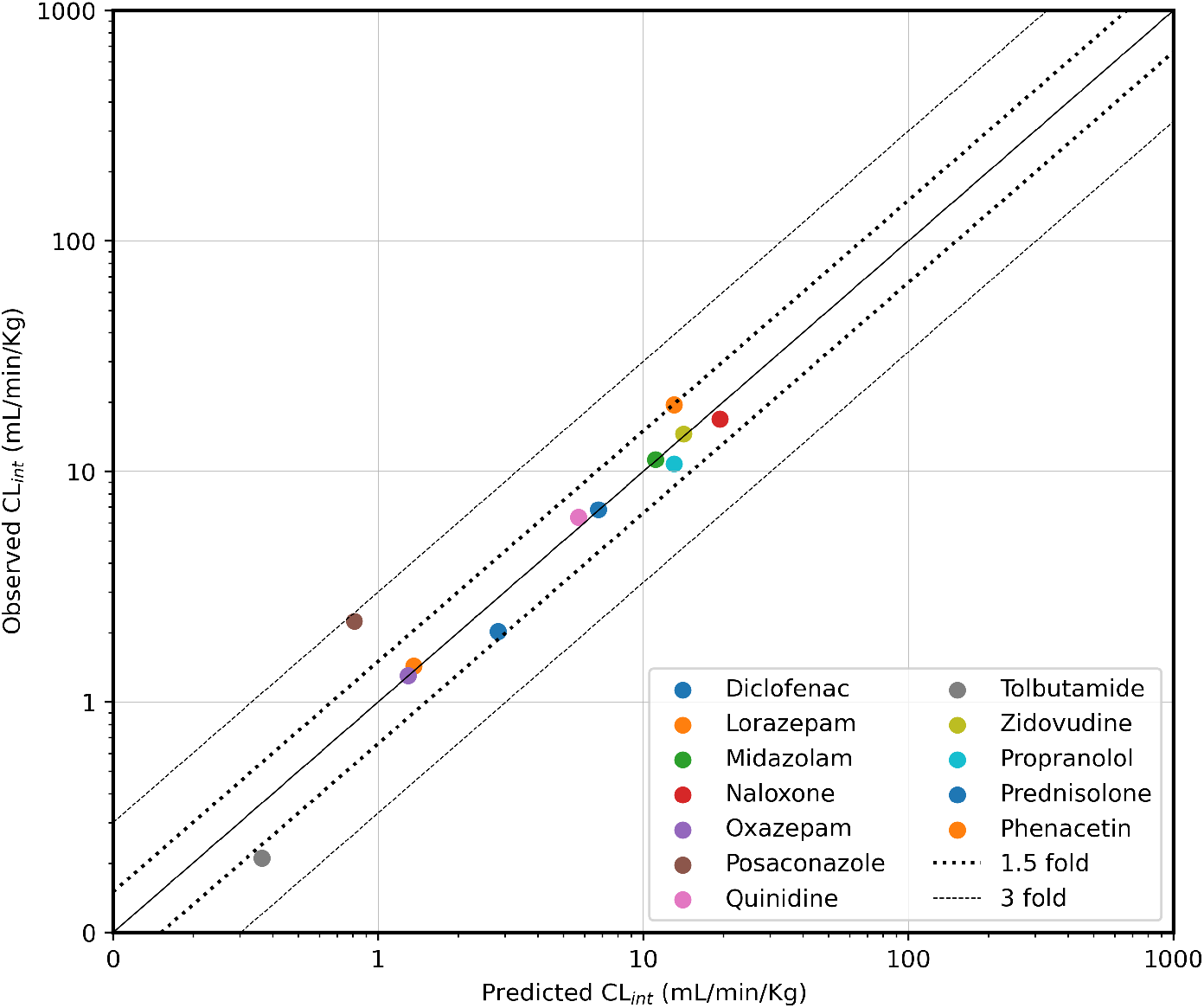
Correlation between observed and predicted *in vivo* intrinsic clearance (CLint) using three-compartment ODE liver chip for 12 drugs from (Docci et al. 2022; Tsamandouras et al. 2017). The solid line shows the line of unity, while dotted line is 1.5-fold and dashed line is 3-fold deviation.

### 3.4 Translation to Human PK

We assessed the impact of accurately predicting human clearance values based on *in vitro* cell-based assays on predicting human PK, using propranolol as a proof-of-concept case study. First, a human PBPK model describing the human kinetics of propranolol was implemented in PK-Sim and qualified with clinical observations. Next, the predicted human clearance value using either the state-of-the-art modelling approach or based on the same on-chip kinetic data, was implemented in the human PBPK model simulating the kinetics after a single oral dose. Further, a population of *n* = 1000 patients was simulated to account for inter-patient variability. As shown in Fig 5, the implemented human PBPK model describes observed clinical data well (using clinical clearance values). When substituting only the clearance value with the state-of-the-art or the digital twin-based values, the impact on simulating human PK becomes apparent, while the standard approach would overpredict (i.e., the on-chip clearance is underpredicted) the human PK (3-fold C_max_, up to 6-fold overprediction of AUC). Moreover, this approach would actually simulate non-negligible concentrations of propranolol left over after 24 h. For repeated daily dosing, this would result in accumulation of propranolol in this hypothetical setting, which would have immediate implications for potential toxicity or efficacy considerations. On the other hand, the digital-twin based approach still slightly overpredicts the AUC and C_max_, however only by 1.5-fold and captures the terminal phase correctly.

## 4. Discussion

The aim of this work is to improve the current prediction of human clearance values and to present a framework for translating *in vitro* findings to relevant clinical situations. The presented integrated translational approach combined quantitative OoC and cell-based assay compound depletion kinetics with an OoC-digital twin to simulate drug kinetics in humans.

Initial investigations revealed the potential to describe clinical clearance values more appropriately than is currently possible with the state-of-the-art approach. This simpler approach lumps biological processes together into a single process – clearance – and uses only minimal information available, e.g., only the cell number and media volume. While biological systems have evolved rapidly in the last decade, especially in the field of organ-on-chip and microphysiological systems, the applied mathematical models to analyse the quantitative complex biological data have been the same for decades (early concept of clearance was introduced by Möllers in 1928, while well-stirred model is 1971).

In contrast, the developed digital twin approach for the organ-on-chip and 3D spheroids comprises three building blocks: biological, hardware, and physicochemical information. The distinction between active and passive processes is achieved by an explicit description of uptake, distribution, and metabolism involved in the biological processes. Further, the digital twin links the architecture of the hardware chip with an advanced mapping of the underlying biology (intracellular compartment). The on-chip kinetics for 32 compounds (six compounds were removed from the initial set due to missing information) was well described, highlighting the drug-specific effects on cellular uptake and hence metabolism. Note that this analysis used the same biological information as used in the state-of-the-art approach, no additional biological experiments were needed or performed to improve the outcome of the digital twin approach.

The predictive power of organ-on-chip and 3D spheroids over conventional approaches was revealed when the depletion data was analysed with the digital twins (Fig 5). Not only was the systematic underprediction issue resolved, but the uncertainty in prediction was also reduced by a factor of 3 (comparing CVs).

Lastly, we aimed to demonstrate the clinical impact of this approach by translating the results to humans using propranolol as a proof-of-concept example. Here, the head-to-head comparison clearly demonstrated the superior power of both quantitative biological data from OoCs and digital twins over state-of-the-art approaches in predicting human PK more appropriately (Fig 7). Although only one compound was used to demonstrate clinical impact, the workflow and process is laid out and easily applicable to other compounds. To the best of our knowledge, this study is the biggest comprehensive report to systematically assess the predictive power of organ-on-chip in the context of use of liver clearance.

**Fig 7.**
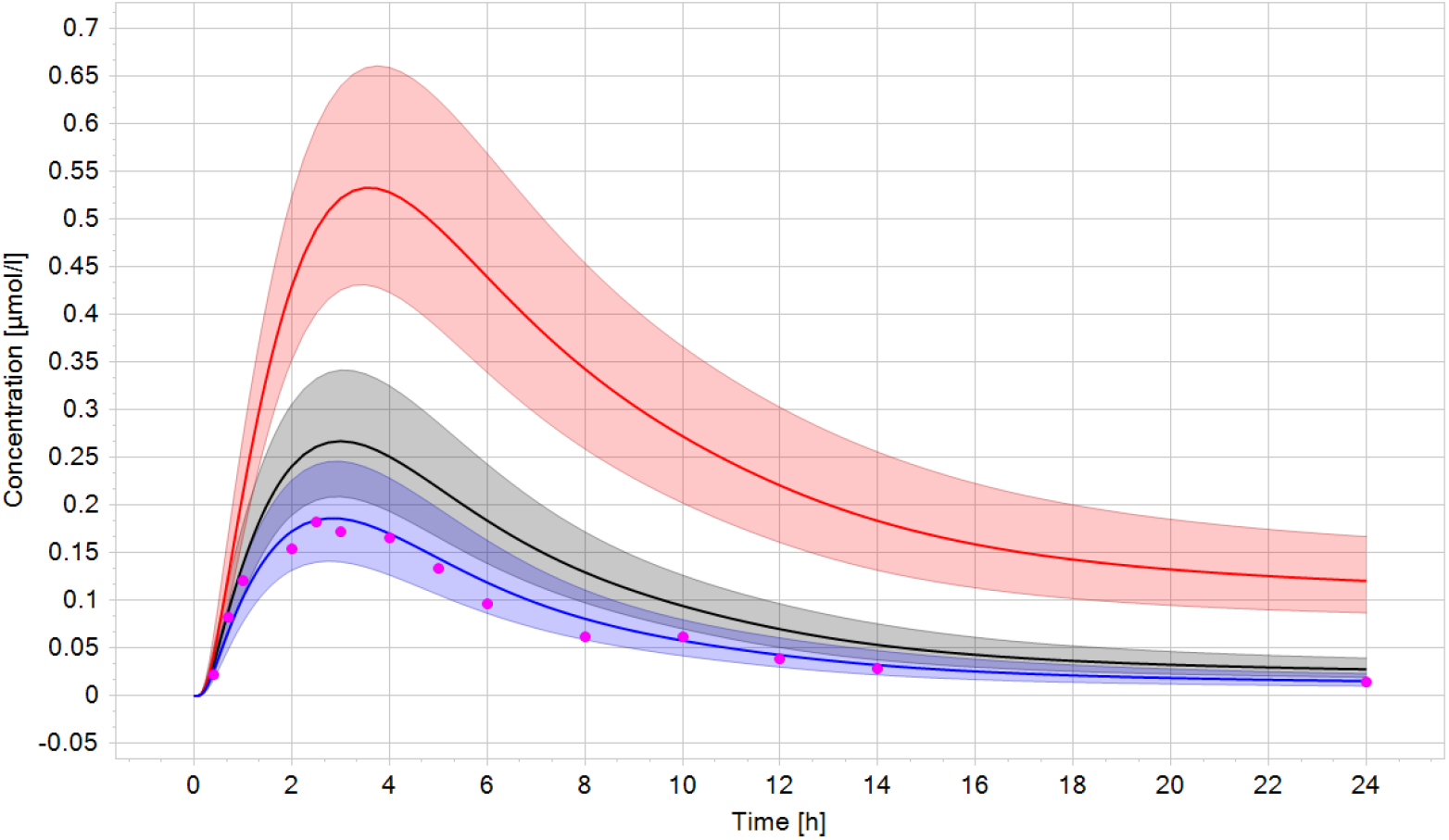
Simulated kinetics of propranolol after a single oral dose (80 mg). Pink dots are clinical observation (digitised from Borgström et al. [25]), while the blue solid line represents the mean of the patient population using the clinical observed clearance value. When using the state-of-the-art based clearance value (red line), the area-under-the-curve is 6-fold overpredicted. In contrast, using the digital twin-based clearance, the AUC is only 1.5-fold overpredicted, also simulating the right kinetics at 24 h (black curve). Shaded areas represent ± 1 SD.

The mathematical algorithm to determine liver clearance depends on the time-concentration profiles and, if available, on intracellular or cell-associated compound levels. The algorithm minimises a cost function by identifying a clearance value such that model prediction and data observation match. The cost function takes both the PK profile from cell culture media and the cell-associated levels into account, which is not the case for the state-of-the-art approach. Further, binding of the compound to plastic/hardware of the chips, to proteins contained in the cell culture media, or any other intracellular lipids can be accounted for to accurately determine liver clearance. DigiLocs, further, also does not depend on a scaling factor, which overcomes the systematic underprediction of state-of-the-art approaches (5-10 fold on average across multiple studies).

So far, limited information is available from the literature or in-house measurements on the observed partitioning of compounds into the intracellular or cell-associated milieu of hepatocytes. If that data becomes available, it may be incorporated compared to the adjustments made in the software to match the clinical clearance values. If the predicted and observed Kp_uu_ values match, the digital twin approach truly improves the prediction. If there is a discrepancy between these values, the fitting process can be re-run including the observed Kp_uu_ value. This would inform the maximum capacity of the system to metabolise the compound. If this final rate is still lower than the observed clinical clearance value two options are possible to understand and improve the prediction:

1. Calculate a correction factor, which is compound-specific and chip-specific and not generic like in the state-of-the-art approaches.
2. Investigate other model-specific parameters to optimise, e.g., permeability or partitioning.

Although initially developed for hepatic clearance, the mathematical model can be employed for toxicity or efficacy-related questions depending on the context of use. In such a setting, time-concentration profiles will be simulated and linked to other, measured biomarkers (e.g., ATP (adenosine triphosphate), TEER (barrier integrity)) to determine IC50 or EC50 values, and parameters to assess toxicity and efficacy, respectively. Likewise, the same integration of complex biological processes, hardware-, and drug-specific information can be used to model other cell and chip types, e.g., a blood-brain-barrier-chip, which is used to determine the permeability of compounds across the barrier.

Eventually, DigiLoCs shall act as decision-support tool for (pharmaceutical) research in estimating the first-in-human doses, assessing human PK, and more importantly, reducing animal experimentation, making drug development efficient, faster, and sustainable.

## 5. Conclusion

The development of digital twins for organ-on-chips, reported here, incorporating advanced mathematical equations and leveraging published data, holds great potential to enhance our understanding of drug behaviour and clinical outcomes. The *in vitro* liver clearance for 32 drugs was predicted using DigiLoCs and a proof-of-concept (translation to human pharmacokinetics) study on propranolol was done. DigiLoCs are envisioned to serve as a decision-support tool for pharmaceutical research, aiding in estimating first-in-human doses, evaluating human pharmacokinetics, and importantly, diminishing reliance on animal experimentation, thereby fostering more efficient, expedited, and sustainable drug development processes. Our approach is generalisable across various physiological contexts and not limited to liver metabolism but may be extended to other organs as well, such as gut metabolism and barrier models such as the brain or placenta.

## Author contributions

Participated in research design: Aravindakshan, Maass. Performed data analysis: Aravindakshan, Maass. Investigation and Methodology: Aravindakshan, Mandal, Pothen, Maass. Writing – Original Draft Preparation: Aravindakshan, Maass. Writing – Review & Editing: Aravindakshan, Mandal, Pothen, Maass.

